# CRISPR/Cas9-mediated genome editing of *Schistosoma mansoni* acetylcholinesterase

**DOI:** 10.1101/2020.07.08.190694

**Authors:** Hong You, Johannes U. Mayer, Rebecca L. Johnston, Haran Sivakumaran, Shiwanthi Ranasinghe, Vanessa Rivera, Olga Kondrashova, Lambros T. Koufariotis, Xiaofeng Du, Patrick Driguez, Juliet D. French, Nicola Waddell, Mary G. Duke, Wannaporn Ittiprasert, Victoria H. Mann, Paul J. Brindley, Malcolm K. Jones, Donald P. McManus

## Abstract

CRISPR/Cas9-mediated genome editing shows cogent potential for the genetic modification of helminth parasites. Here we report successful gene knock-in (KI) into the genome of the egg of *Schistosoma mansoni* by combining CRISPR/Cas9 with single-stranded oligodeoxynucleotides (ssODNs). We edited the acetylcholinesterase (AChE) gene of *S. mansoni* targeting two guide RNAs (gRNAs), X5 and X7, located on exon 5 and exon 7 of Smp_154600, respectively. A CRISPR/Cas9-vector encoding gRNA X5 or X7 was introduced by electroporation into eggs recovered from livers of experimentally infected mice. Simultaneously, eggs were transfected with a ssODN donor encoding a stop codon in all six frames, flanked by 50 nt-long 5’- and 3’-homology arms matching the predicted Cas9-catalyzed double stranded break at X5 or X7. Next generation sequencing analysis of reads of amplicon libraries spanning targeted regions revealed that the major modifications induced by CRISPR/Cas9 in the eggs were generated by homology directed repair (HDR). Furthermore, soluble egg antigen from AChE-edited eggs exhibited markedly reduced AChE activity, indicative that programmed Cas9 cleavage mutated the AChE gene. Following injection of AChE-edited schistosome eggs into the tail veins of mice, a significant decrease in circumoval granuloma size was observed in the lungs of the mice. Notably, there was an enhanced Th2 response involving IL-4, −5, −10, and-13 induced by lung cells and splenocytes in mice injected with X5-KI eggs in comparison to control mice injected with unmutated eggs. A Th2-predominant response, with increased levels of IL-4, −13 and GATA3, also was induced by X5 KI eggs in small intestine-draining mesenteric lymph node cells when the gene-edited eggs were introduced into the subserosa of the ileum of the mice. These findings confirmed the potential and the utility of CRISPR/Cas9-mediated genome editing for functional genomics in schistosomes.

**Author Summary:** Schistosomiasis is the most devastating of the parasitic helminth diseases. Currently, no vaccines are available for human use and praziquantel is the only available treatment raising considerable concern that drug resistance will develop. A major challenge faced by the schistosomiasis research community is the lack of suitable tools to effectively characterise schistosome gene products as potential new drug and/or vaccine targets. We introduced CRISPR/Cas9 mediated editing into *S. mansoni* eggs targeting the gene encoding acetylcholinesterase (AChE), a recognized anthelminthic drug target. We found that the major modifications induced by CRISPR/Cas9 in the eggs were generated by homology directed repair (HDR). This platform provides a unique opportunity to generate precise loss-of-function insertions into the schistosome genome. We pre-screened the activity of two guide RNAs of the AChE gene and compared/validated the mutation efficacy using next-generation sequencing analysis at the genomic level and phenotypic modifications at the protein level. That resulted in reduced AChE activity observed in AChE-edited eggs, and decreased lung circumoval granuloma size in mice injected with those edited eggs. The CRISPR/Cas9-genome editing system we established in this study provides a pivotal platform for gene functional studies to identify and test new anti-schistosome intervention targets, which can be extended to the other human schistosome species and other important parasitic helminths.

## Introduction

Schistosomiasis remains one of the most prevalent, chronic and insidious of the tropical parasitic diseases. Human infection begins when larval schistosome cercariae penetrate the skin. Once in the bloodstream, the cercariae transform into schistosomula which migrate to venules associated with specific organs (e.g. liver, intestine, bladder) where they mature as adult male and female worms and rapidly start to reproduce. Their eggs release molecules that damage host tissues through induction of extreme granulocytic responses and tissue scarring. No effective schistosomiasis vaccine is available and treatment is entirely dependent on a single drug, praziquantel (PZQ) [1]. Despite the wide-spread use of PZQ, with over 40 years of mass drug administration programs, the number of people infected, particularly in Africa, has not decreased substantially and, in fact, may exceed published figures [2]. Furthermore, this MDA strategy is unsustainable long term and the spectre of the likely generation of PZQ-resistant parasites is a constant threat. Complete genomic sequences are available for *Schistosoma japonicum* [3], *S. mansoni* [4] and *S. haematobium* [5]. The complex, multi-generational life cycles of schistosomes and their recalcitrance for genetic manipulation has resulted in major difficulties for gene editing [6]. Notably, the paucity of molecular tools to manipulate schistosome gene expression has markedly limited our ability to define key metabolic pathways, thereby hampering discovery of new anti-schistosome drugs or vaccines.

CRISPR/Cas9 is an advanced genome editing tool and RNA-guided system whereby a 20-base guide RNA (gRNA) directs the Cas9 nuclease from *Streptococcus pyogenes* to cleave a target gene, generally providing high specificity and minimal off-target site effects. The endonuclease makes a double-strand break (DSB) at the target site [7]. Endogenous DSBs in eukaryotes are repaired by at least two repair mechanisms [8]: non-homologous end joining (NHEJ), which results in nucleotide insertions and deletions [9]; and homology directed repair (HDR) which utilizes a repair template. In terms of heterogeneity of indels introduced at Cas9-induced DSBs, allelic editing frequencies are variable [10]. The prokaryotic CRISPR/Cas9 system was first adapted for *Caenorhabditis elegans* in 2013 [11] and used to generate transgenic organisms and introduce site-specific, heritable mutations [12]. It was further extended for genome editing in diverse organisms, including human protozoan parasites *Toxoplasma gondii, Plasmodium falciparum, Trypanosoma cruzi* and *Leishmania* species [13]. Deployment of the CRISPR/Cas9 system for genome editing [14] extends the range of experimental approaches to interrogate the host-parasite relationship and test novel vaccines or drugs. Post-transcriptional gene silencing (RNA interference), which has been developed over the last 15 years for loss-of-function research in schistosomes, is transient and limited in maintaining inheritance of the targeted gene. Ittiprasert *et al* pioneered the CRISPR/Cas9 editing system in *S. mansoni* eggs, targeting a single gRNA of omega-1 (*ω1*), which plays a crucial role in Th2 polarization and granuloma formation [15]. Sankaranarayanan *et al* compared CRISPR/Cas9 mediated deletions in different life cycle stages of *S. mansoni* including adult worms, sporocysts and eggs, targeting the gene *SULT-OR* which is involved in oxamniquine resistance [16] (preprint data). Arunsan *et al*. also undertook a similar approach to modify the granulin (*Ov*-GRN-1) gene in the liver fluke *Opisthorchis viverrini* which plays a key role in virulence morbidity during opisthorchiasis [17]. These pivotal studies demonstrated that programmed genome editing is feasible in flatworm parasites.

Effective site-specific gene modification and phenotyping will drive innovation and a better understanding of schistosome pathogenesis, biology and evolution [6]. To expand functional genomic investigations of schistosomes and, in particular, to advance our understanding of the function of acetylcholinesterase (AChE) in these blood flukes, we undertook CRISPR/Cas9-mediated knock-out (KO) and knock-in (KI) in adult *S. mansoni* and eggs targeting this key enzyme. AChE is the target of a number of currently approved and marketed anthelmintics [18-20]. Furthermore, AChE plays an important role in the adult schistosome neuromusculature cholinergic system [21, 22], and is intimately involved in muscle function [23, 24] and other essential activities such as feeding, sexual maturation and mating in mature worms [24]. We previously demonstrated that AChE is present on the external tegumental membrane and in the musculature of adult schistosomes [21, 25]. AChE also occurs within the eggs and in the host cellular components of the advanced granulomas around the parasite eggs entrapped in various organs [26] indicating an involvement in granuloma formation as a result of its role in inhibiting the host IL-4 response [26].

Here, we used programmed genome editing to modify the AChE-encoding locus in the genome of the eggs and adult developmental stages of *S. mansoni* using CRISPR/Cas9-based KO and KI approaches. We determined the efficiency of CRISPR/Cas9-mediated midifications by next generation sequencing analysis (NGS) on genomic DNA extracted from mutated parasites. We compared KO/KI efficacy by targeting two gRNAs of AChE distanced with 14.3kb genomic sequence individually and in combination. In addition, we assessed *in vivo* immune responses induced by AChE-KI eggs in lung cells, splenocytes and small intestine-draining mesenteric lymph node cells following injection of AChE KI eggs into the tail vein of mice or into the small intestinal subserosa of mice.

## Results

### Design of two single guide RNAs

Two guide RNAs were designed to target residues 722 – 741 in exon 5 (X5) and 1738 – 1757 in exon 7 (X7), respectively, of the acetylcholinesterase gene (AChE, Smp_154600, Uniprot Q71SU7_SCHMA), which is located on *S. mansoni* chromosome 1 (**Fig 1A**). The nucleotide residues complementary to guide RNA-X5 and -X7 are adjacent to the protospacer adjacent motifs (PAM), TGG and AGG, respectively, with the predicted Cas9 cleavage site located three residues upstream of the PAM. These PAMs (TGG or AGG) and the nucleotide sequences complementary to these two gRNAs are highly specific and are absent from two paralogues of Smp_154600, i.e., Smp_136690 and Smp_125350. These two paralogues share 34-38% nucleotide identity and 25-26% amino acid identity to Smp_154600. The Smp_136690 and Smp_125350 genes exhibit a distinct exon/intron structures (**S1 Fig**) (although this may reflect imprecise annotation of the draft genome sequence). **S2 Fig** presents the amino acid sequence alignment and predicted functional motifs of Smp_154600, Smp_136690 and Smp_125350.

**Fig 1.**
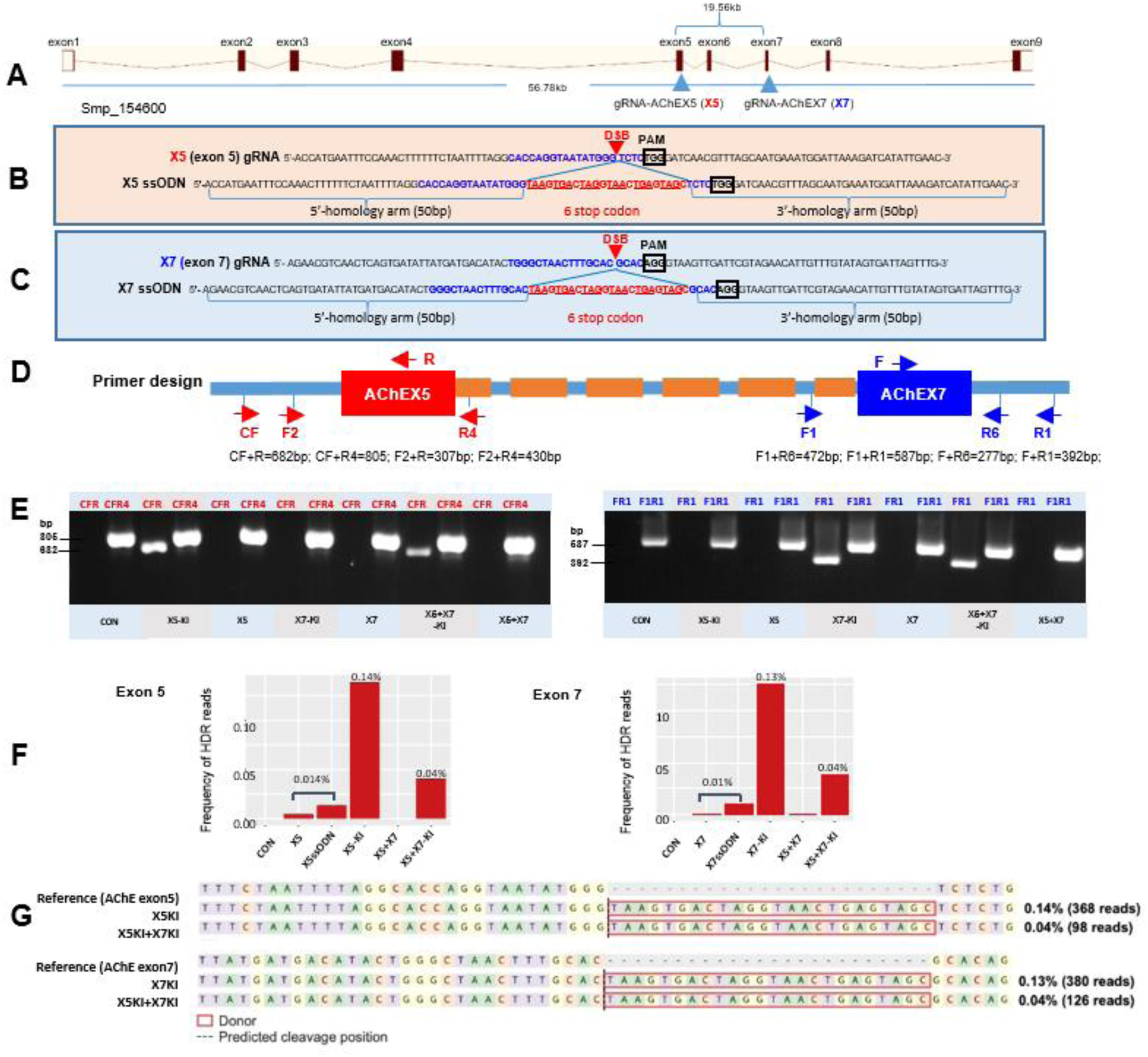
CRISPR/Cas9-mediated editing of the acetylcholinesterase (AChE) locus. (A) Schematic diagram of the AChE (Smp_154600) gene in *S. mansoni*. Two guide RNAs (gRNAs) were designed to target AChE: X5 targeting exon 5 and X7 targeting exon 7. Partial nucleotide sequence of (B) AChE exon 5 and (C) exon 7 indicating the respective gRNA sequence, target site, predicted double stranded break (DSB), protospacer adjacent motif (PAM), and sequence of the single-stranded oligodeoxynucleotide (ssODN) donor template. Homology arms of 50 nt flank a central 24 nt six-stop-codon transgene. (D) Schematic diagram of AChE indicating positions and directionality of primer binding sites (arrows). Primers in red target AChE exon 5 (CF, F2, R, R4) and primers in blue target AChE exon 7 (F1, F, R6, R1). The control PCR amplicons were generated using the CF+R4 primers (red arrows) targeting the fragment around gRNA-X5, and F1+R1 primers (blue arrows) targeting the fragment around gRNA-X7. (E) PCR products demonstrating CRISPR/Cas9-mediated editing in exon 5 (left panel) and 7 (right panel) of the AChE gene. Genomic DNA extracted from eggs treated with CON, X5-KI, X5, X7-KI, X7, X5+X7-KI and X5+X7 were used as the PCR template. Evidence of X5-KI and X7-KI revealed by amplicons of the expected sizes in lanes CFR (682 bp) and FR1 (392 bp), respectively, spanned the mutated site in the genomic DNAs pooled from schistosome eggs, including positive controls flanking the insert site of gRNA-X5 (CFR4, 805 bp) and the insert site of gRNA-X7 (F1R1, 587 bp). The control DNA result shown in this gel was isolated from eggs electroporated with negative control vector (CON) only. (F) Frequency of homology directed repair (HDR) at AChE exon 5 (left panel) and exon 7 (right panel) in amplicon deep sequencing data. Amplicons at exon 5 were generated using primer pair F2+R4, and amplicons at exon 7 were generated using primer pair F1+R6 as indicated in (D). HDR reads were detected using CRISPResso2 and confirmed using fasta36. The number of confirmed HDR reads are shown, expressed as a percentage of aligned reads. (G) Sequence alignments and HDR frequencies for amplicon deep sequencing data. Nucleotide sequences for AChE exon 5 (upper panel) and exon 7 (lower panel), extending 30 bp either side of the respective predicted cleavage position, are shown. At both loci, the presence of a 24 nt transgene (red outline) was confirmed. HDR frequencies, expressed as a percentage of aligned reads, are shown to the right of each KI group. Plots were based on default output from CRISPResso2.

### Site-specific integration of exogenous DNA confirms CRISPR/Cas9 activity in schistosomes

We investigated the activity and efficiency of programmed gene knockout (KO) of the AChE gene (Smp_154600) of *S. mansoni*. The two AChE locus specific gRNAs (X5 and X7 described above) were cloned into the GeneArt CRISPR Vector, which encodes Cas9 of *Streptococcus pyogenes* driven by the human cytomegalovirus (CMV) promoter and gRNA (X5 or X7) driven by the human U6 promoter. Freshly perfused *S. mansoni* adult worms and eggs isolated from the livers of infected mice were transfected with the plasmid vector constructs by square wave electroporation.

Homology directed repair (HDR) of CRISPR/Cas9-induced double stranded breaks (DSBs) at a gene locus in the presence of donor DNA template was reported in the human threadworm *Strongyloides stercoralis* [27]. Accordingly, as a template for HDR of chromosomal DSBs, a single stranded oligodeoxynucleotide (ssODN) of 124 nt in length targeting X5 (X5ssODN, **Fig 1B**) was delivered with the X5-CRISPR vector (X5 knock-in, X5-KI) and ssODN targeting X7 (X7ssODN, **Fig 1C**) was delivered with X7-CRISPR vector (X7-KI); both X5ssODN and X7ssODN were delivered with X5-CRISPR vector and X7-CRISPR vector (X5+X7-KI) into eggs and adult worms.

Both X5ssODN and X7ssODN include a short transgene encoding six stop codons flanked by 5’- and 3’-homology arms, with each arm 50 nt in length complementary to the genome sequence of exon 5 and 7, respectively, on both the 5’ and 3’ sides of the programmed Cas9 cleavage site (**Fig 1B, 1C**). Given that the donor ssODN includes a short transgene in order to facilitate genotyping, PCR was performed using template genomic DNA extracted from the CRISPR/Cas9-treated eggs and adults to detect the extent of KI. Targeting AChE gRNA X5, a reverse primer termed R, specific for the stop codon region of the donor ssODNs, was paired with two discrete forward primers (termed CF and F2, **Fig 1D**); R4 was designed as a reverse primer to pair with CF and F2 as positive (wild type) control (**Fig 1D**). Targeting gRNA X7, a forward primer termed F, specific for the stop codon region of the donor ssODN transgene, was paired with two discrete reverse primers (termed R1 and R6, **Fig 1D**); F1 was designed as forward primer to pair with R1 and R6 as positive control (**Fig 1D**). S1 Table provides the nucleotide sequences of these primers.

At AChE X5 amplicons of the expected sizes for primer pairs CF+R (682 bp) and F2+R (307 bp) were observed in eggs (**Fig 1E**) and adults (**S3A Fig**) in genome-edited knock in (KI) groups X5KI and X5+X7-KI, but not in group X7-KI, indicating successful insertion of the ssODN transgene and resolution of the DSB. PCR results from the control groups, namely parasites electroporated with negative control CRISPR vector (CON) or media, did not show bands at 682 bp or 307 bp. AChE X7 amplicons of 392 bp (primers F+R1) were observed in groups X7-KI and X5+X7-KI in both eggs (**Fig 1E**) and adults (**S3 Fig**), but not in group X5-KI and the CON groups (**Fig 1E**), indicating programmed insertion of the ssODN transgene. The PCR profiles obtained for groups of parasites treated with ssODN only (including X5ssODN and X7ssODN) were the same as the control groups. Importantly, Sanger sequencing analysis of the KI amplicons confirmed the insertion of the transgene into the AChE locus at X5 and X7 at the predicted cleavage sites.

### Viability of eggs and adult worms after electroporation

The viability of eggs isolated from livers of mice infected with *S. mansoni* cercariae and post-electroporation revealed hatching rates of 70-80% (**S4 Fig**). The hatching rate was slighted decreased, but not significantly (*p*=0.16), in eggs subjected to electroporation compared with those that were not (**S4 Fig**), indicating that electroporation did not markedly affect egg viability. Microscopy confirmed adult worms of *S. mansoni*, cultured for two days post-electroporation, were alive.

### Programmed modifications at exons 5 and 7 of the AChE locus within the genome of the eggs of *S. mansoni*

To characterize and quantify the modifications at the nucleotide level resulting from CRISPR/Cas9-induced AChE-KO/I into the schistosome egg, we used an amplicon NGS approach, with specific forward primers (F2 for X5 and F1 for X7, **Fig. 1D**) and reverse primers (R4 for X5 and R6 for X7, **Fig. 1D**) designed to target the AChE locus in regions flanking the predicted programmed modifications. Barcoded amplicon libraries were constructed from pooled genomic DNA of independent groups of eggs with respect to the several guide RNAs and donor ssODN combinations. The sequencing was performed on the Illumina MiSeq platform. CRISPResso2 [28] was used to analyse deep-coverage sequence reads. On average, 312,000 reads per sample aligned to the respective reference amplicon sequence of the Smp_154600 locus (S2 Table).

### Modifications observed at AChE exon 5

We used the CRISPResso2 software to analyze the sequenced amplicon reads obtained using the primers F2 and R4 targeting exon 5 of AChE. After filtering the reads based on the F2+R4 primers, 0.21% of reads from eggs subjected to X5-KI treatment exhibited sequence variations ascribable to chromosomal DSB repair by NHEJ or HDR. By running CRISPResso2 with an increasing window size parameter (value of 1, 20, 100, and 0) which defines the size (in bp) of the quantification window extending from the DSB, we found that the amount of NHEJ modifications, particularly NHEJ substitutions, also stably increased across both the negative control and experimental samples (see **S5 Fig**). Therefore, we kept the quantification window size around each gRNA to the default value of 1 to limit the amount of PCR and/or sequencing errors from being inappropriately quantified as modified reads and we found that the majority (>95%) of modified reads with NHEJ were due to substitutions, and not insertions or deletions (see **S2 Table**). Of the reads that were modified in the samples treated with X5-KI, 0.14% were confirmed by fasta36 to be HDR modified reads (**Fig 1F**). We also analysed samples treated with X5+X7-KI and found 0.04% of reads were confirmed HDR events, which is lower than the percentage of HDR reads in samples treated only with X5-KI (**Fig 1F**). Additionally, we detected a small number of HDR events in the negative control samples which may have been due to sample cross-contamination possibly occurring during barcoding in the amplicon library preparation. The level of background noise for HDR detection was set to 0.014%, since the negative control samples contained HDR events below this value. By subtracting the level of background noise for HDR detection at exon 5 from the rate of confirmed HDR in X5-KI (0.14%) and X5+X7-KI (0.04%), we estimated that 0.12% and 0.03% of reads in the X5 experimental samples carried CRISPR/Cas9-induced HDR events, respectively.

### Modifications at AChE exon 7

We used CRISPResso2 to analyse the raw reads from samples that were amplified using the F1+R6 primers targeting AChE exon 7. After filtering reads based on primers F1+R6, 0.26% of reads from the sample treated with X7-KI were found modified by NHEJ or HDR: 0.13% were confirmed as HDR events by fasta36 analysis, while the other 0.13% were NHEJ modifications, majority of which were substitutions and not insertions or deletions (see **S2 Table**). Similar to the findings with exon 5, the samples treated with X5+X7-KI contained fewer confirmed HDR events (0.04%) than samples treated with the single guide and donor combination (X7-KI). For exon 7, the level of background noise for HDR detection was set to 0.0105%, since the negative control (X7 and X7ssODN groups) samples contained HDR events below this value. By subtracting the level of background noise for HDR detection at exon 7 from the rate of confirmed HDR in the experimental samples, we estimated that 0.1195% and 0.0295% of reads represented CRISPR/Cas9-induced HDR events in the X7 treatment samples, respectively.

### Decreased AChE activity in CRISPR/Cas9-mediated KI eggs and adult worms

The AChE activities of soluble egg antigen (SEA) extracted from eggs, and soluble worm antigen preparation (SWAP) obtained from adult worms of *S. mansoni*, were determined following CRISPR/Cas9-mediated AChE-KI. Significant decreases in AChE activity were observed in SEA (**Fig 2A**) extracted from eggs treated with X5-KI (10.7%, *p*=0.001), X7-KI (8.3%, *p*=.006) and X5+X7-KI (13.4%, *p*=0.0002) compared with SEA from control eggs electroporated with CON. Remarkable reductions in AChE activity were also detected in SWAP (**S3B Fig**) extracted from adults treated with X5-KI (21%, *p*=0.023), X7-KI (11.8%, ns) and X5+X7-KI (23.8%, *p*=0.012) compared with control adults electroporated with CON. The control groups, including SEA (or SWAP) extracted from wild type parasites and parasites electroporated with media, CON or ssODN only, exhibited a similar level of AChE activity, and no significant differences were observed amongst them.

**Fig 2.**
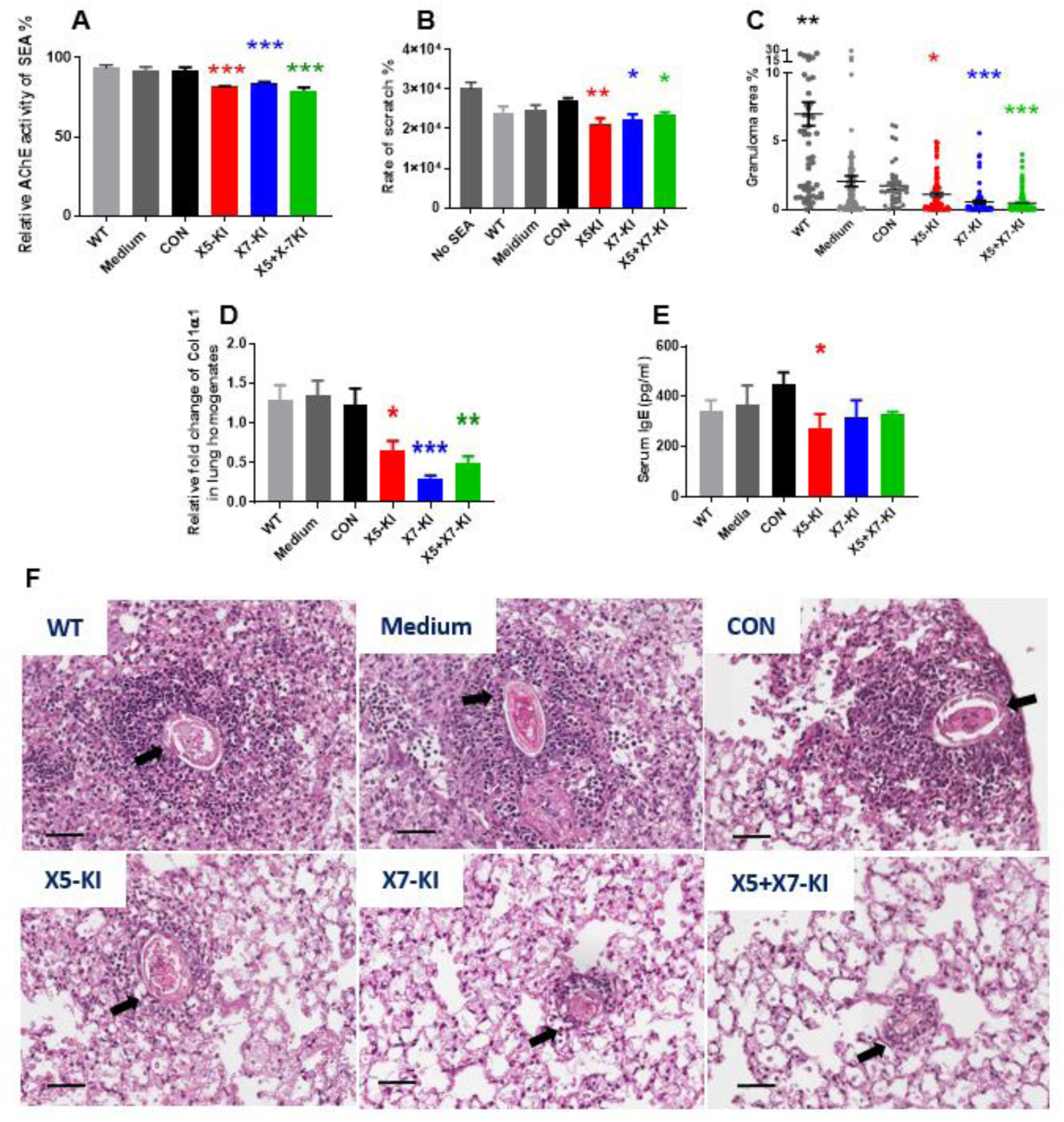
Effects of AChE-KI eggs of *S. mansoni* on AChE enzyme activity, LX-2 cell growth and granulomatous inflammation, collagen expression and serum IgE levels in mice. (A) Diminished AChE activity of SEA extracted from *S. mansoni* eggs after incubation with medium, CON, X5-KI, X5, X7-KI, X7 and X5-KI+X7-KI, respectively. SEA extracted from Wild type (WT) eggs were used as positive control. (B) Monitoring LX-2 cell growth on day 3 after being incubated with mutated/unmutated SEA. The scratch wound assay using the IncuCyte system measured the migration of LX-2 cells. The total area (μm^2^) was obtained at every time point (from day 0 to day 3) until wound closure, and triplicate measures were compared to calculate an average closure rate per group (area of original wound/time to closure in hours). No SEA: cells cultured in medium without SEA. (C) Effect of AChE-KI eggs on granuloma formation in the lungs of mice i.v. injected with eggs treated with X5-KI, X7-KI and X5-KI+X7-KI, respectively. A control group of mice received untreated WT eggs or CON eggs. All mice were euthanized 2 weeks after injection. Granuloma sizes were determined as the ratio of the granuloma area to the egg area. The scatter plot shows the mean ± SE. of the data pool representing all granulomas from all lungs from each experimental group (WT: n=60; CON=40; X5-KI: n=88; X7-KI: n=72; X5+X7-KI: n=120); each group included 10 mice. (D) Transcription level of collagen type 1 (alpha 1-COL1α1) in *S. mansoni* eggs in different experimental groups. HPRT (hypoxanthine guanine phosphoribosyl transferase) was used as a reference gene to normalise data [87]. (E) Total serum IgE levels in mice two weeks following i.v. injection of WT eggs of *S. mansoni* and eggs treated with medium, CON, X5-KI and X7-KI, X5+X7-KI. For panels A-E, each experiment was performed in duplicate and all data are presented as the mean ± SE. One-way ANOVA analysis was used to establish statistical significance compared with the CON control: * = *p* value≤0.05, ** = *p* value≤0.001, *** = *p* value≤0.0001. (F) Granuloma size in lung sections from mice i.v. injected with eggs of *S. mansoni* in different groups, examined microscopically following H & E staining. Scale bars, 50µm. WT, wild type parasites; Medium, parasites electroporated with medium; CON, parasites treated with negative control vector.

### Inhibition of migration of LX-2 cells *in vitro* by SEA isolated from AChE-KI eggs

It is recognized that hepatic stellate cells (HSC) resident within liver granulomas are critical players involved in hepatic schistosomiasis being one of the major cell types responsible for the egg-induced immunopathology and resulting fibrosis [29, 30]. To determine further phenotypic changes induced in eggs by AChE-KI we examined the effect of treating LX-2 cells with SEA isolated from AChE-KI eggs, by scratch wound assay. LX-2 is a developed immortalised human HSC cell line that exhibits the key features of primary HSC [31]. Real time monitored cell growth assays using the IncuCyte system were undertaken to test the effects of SEA isolated from AChE-KI and control eggs on the proliferation of LX-2 cells. Cell growth images on days 0, 1 and 2 post-incubation with SEA extracted from mutated/unmutated eggs are shown in **S6 Fig**. The scratch wound assay revealed that, after 3 days culture with SEA (30 µg/ ml), the wound width of LX-2 cells was significantly larger in cells subjected to SEA extracted from eggs treated with X5-KI, X7-KI or X5-KI+X7-KI compared with SEA from eggs electroporated with EV (or medium or WT eggs) (**Fig 2**B). Our analysis showed that the rate of scratch wound closure in SEA-treated cells decreased by 22% in X5-KI SEA (*p*=0.002), 18% in X7-KI SEA (*p*=0.01) and 14% in X5+X7-KI SEA (*p*=0.03) compared with the CON SEA (**Fig 2B**). These data show that SEA derived from CRISPR/Cas9-modified KI eggs targeting either AChE X5 or X7 significantly inhibited HSC migration.

### Decreased granulomatous inflammation and collagen expression in lungs of mice injected with CRISPR/Cas9-mediated AChE-KI eggs

AChE-KI (X5-KI, X7-KI and X5+X7-KI) eggs were injected into the lateral vein of the tails of mice to determine whether AChE-KI eggs affected the development of granulomas associated with pulmonary schistosomiasis *in vivo*. In this model, eggs are transported to the lungs via the bloodstream resulting in subsequent granuloma formation [32]. Two weeks post-injection, mice were euthanized and the lungs of each animal were removed for histological analysis. Representative digital microscopic images of lung tissues acquired through the Aperio ImageScope software are shown in **Fig 2C**. Notably, decreased granuloma area was observed in the AChE-KI (X5-KI, X7-KI and X5+X7-KI) groups (**Fig 2F**). The relative areas of granulomas surrounding individual eggs were quantified and showed 4.3-fold (*p*=0.006), 8.2-fold (*p*<0.0001) and 10-fold (*p*<0.0001) reductions in the X5-KI group, X7-KI and X5+X7-KI group, respectively, compared with the size of granulomas formed around eggs electroporated with CON (**Fig 2C**). These results documented a marked deficiency in the induction of pulmonary granulomas by the injected AChE-KI eggs compared with CON eggs of *S. mansoni*. However, we noted a significantly increased granuloma area detected in the WT group compared with those in the Medium/CON groups (**Fig 2C, 2E**), suggesting that electroporation may have contributed to the reduced egg fitness in the infected mice. During granuloma development in liver, activated hepatic stellate cells deposit a ring of collagen to encapsulate what then becomes the ‘core’ of the granuloma. We thus used portions of the mouse lung to assess the transcription levels of collagen (collagen type 1, alpha 1-COL1α1) in the AChE-KI groups by qPCR (**Fig 2D**). There were significant decreases in collagen mRNA expression in the mice injected with X5-KI eggs (75%, *p*=0.036), X7-KI eggs (46%, *p*<0.0001) and X5+X7-KI eggs (59%, *p*=0.005) compared with control eggs treated with CON only (**Fig 2D**). No statistical difference in collagen mRNA expression was observed in the lungs of mice injected with WT eggs or injected eggs treated with CON (*p*>0.05).

### Serum IgE levels decreased in mice injected intravenously (i.v.) with X5-KI eggs

As a marker of Th2 response polarization in schistosome infection [33], mouse IgE levels were measured in serum samples collected from mice two weeks post-iv-injection with eggs treated with X5-KI, X7-KI or X5+X7-KI. A significant decrease in the level of IgE (*p*=0.04) was observed in mice injected with X5-KI eggs, but not X7-KI or X5+X7-KI eggs, compared with animals injected with CON treated eggs (**Fig 2E**).

### Cytokine responses generated in lung cell isolates and splenocytes of mice injected i.v. with AChE-KI eggs

To further evaluate the immune response generated in mice by AChE-KI eggs, cytokines secreted from lung cells and splenocytes were quantified using LEGEND plex mouse Th1/Th2 panel kits. Both lung cells and splenocytes isolated from individual mice at two weeks post egg-injection were incubated with SEA extracted from normal *S. mansoni* liver eggs for 72 hours. Culture supernatants were then collected for cytokine population analysis. We found that the levels of IL-2, 4, 5, 10, 13 in lung cell (**Fig 3A**) and IL-4, 5, 6, 10, 13 in splenocytes (**Fig 3B**) were significantly increased in mice injected with X5-KI eggs in comparison to control mice injected with CON-eggs. Significantly enhanced levels of IL-6 and TNFα in splenocytes and TNFα in lung cells were observed in mice injected with X7-KI eggs (**Fig 3A**). However, significant change in cytokine response was not evident in the X5+X7-KI egg injected group.

**Fig 3.**
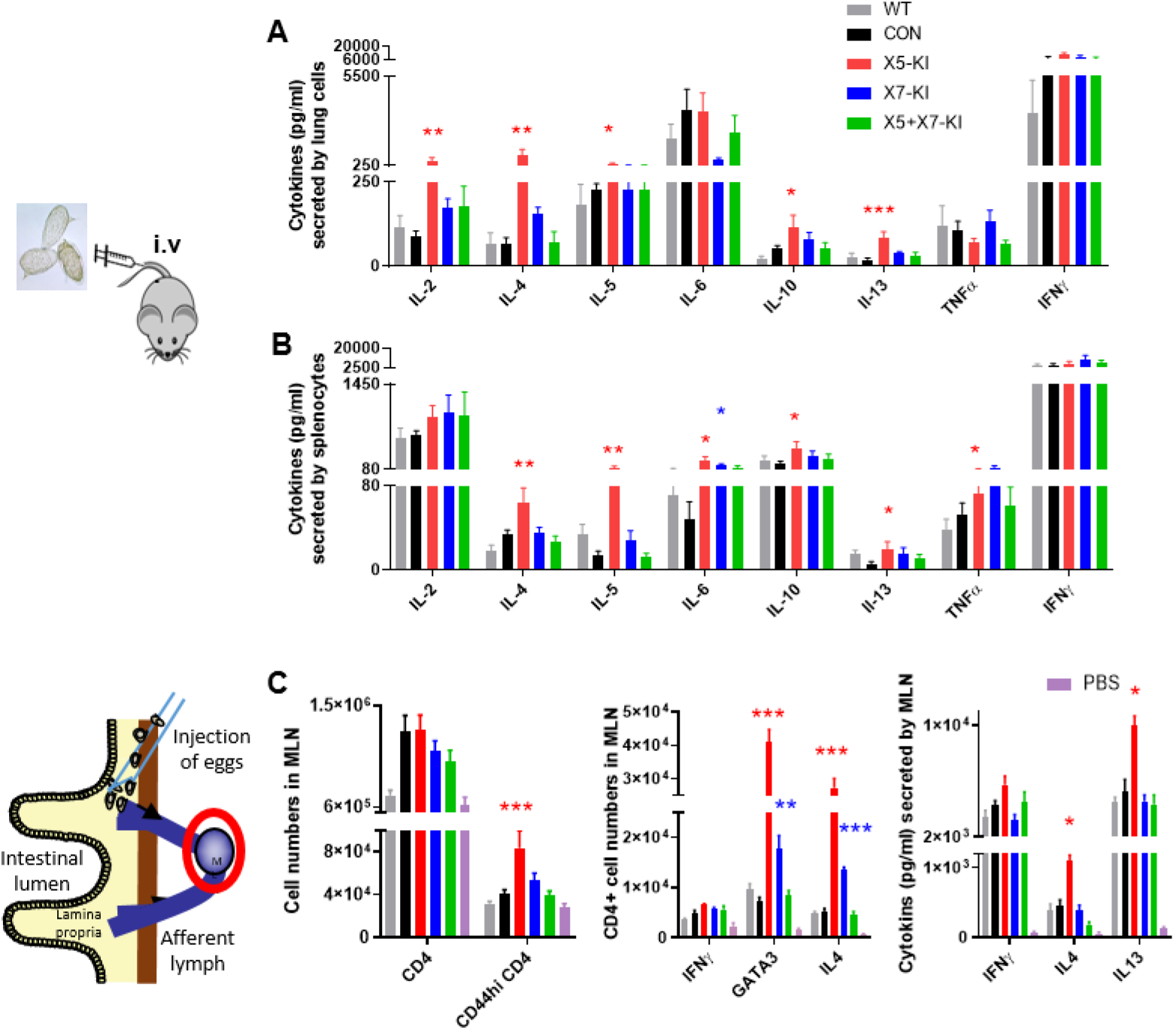
Cytokine production by lung cell suspensions, splenocytes and MLN cells collected from mice injected with AChE-KI eggs. To test cytokine secretion of immune cells derived from lung and spleen, lung cells and splenocytes from mice were analysed at 2 weeks after i.v. injection, via the tail vein, with WT eggs of *S. mansoni* and eggs treated with empty-vector (CON), X5-KI, X7-KI and X5+X7-KI. Cytokines (IFNγ, TNFα, IL-2, IL-4, IL-5, IL-6, IL-10 and IL-13) were quantified in culture supernatants of (A) lung cells and (B) splenocytes restimulated with SEA for 72 hours in the different treated and untreated groups. To determine the cytokine profiles of intestinal immune responses induced by AChE-KI eggs, ileum draining mesenteric lymph node (MLN) cells were isolated from mice on day 5 after subserosal injection with WT and CON eggs of *S. mansoni* and eggs treated X5-KI, X7-KI and X5+X7-KI. (C) Left panel, the proportion of CD4^+^ T cells, CD44hi CD4^+^ cells; Middle panel, the proportion of GATA3-, IFNγ- and IL-4-producing MLN CD4^+^ T cells isolated from MLN cells for each mouse were determined; right panel, secreted cytokines (IFNγ, IL-4 and IL-13) were quantified in culture supernatants following 72 hours culture. WT, Wild Type; CON, Control; IL, interleukin; IFN, interferon; TNFα, Tumor necrosis factor alpha; PBS, Phosphate-Buffered Saline. Each experiment was performed in duplicate; data are presented as the mean ± SE and one-way ANOVA analysis was used to establish statistical significance compared with the CON: \**p*<0.05, \*\**p*<0.01, \*\*\**p*<0.001

### IL-4, IFNγ, Ctla4 and RetnlA mRNA expression in lungs of mice injected i.v. with AChE-KI eggs

To determine whether AChE-KI eggs would change the alternative activation of macrophages, an important hallmark of innate type 2 immunity [34], qPCR was performed on lungs from i.v. injected animals; the expression of IL-4 and IFNγ, Ctla4 (cytotoxic T-lymphocyte-associated protein 4) and RetnlA (resistin-like molecule alpha) were also measured. We found increased levels of IL-4 in mice injected with X5-KI (**S7A Fig**) and enhanced Ctla4 responses were detected in mice injected with X7-KI eggs compared with mice injected with CON-eggs (**S7 Fig**). However, a significant decline in the level of IL-4 was observed in mice injected with CON eggs compared with those injected with WT-eggs (**S7A Fig**). No significant difference was observed in the expression of IFNγ or RetnlA in lungs of mice injected with AChE-KI eggs compared with those receiving CON eggs (**S7B and S7C Fig**).

### Immune responses in mouse mesenteric lymph node (MLN) cells induced by intestinal injection of AChE-KI eggs

To address effects on intestinal immune responses, which have been linked to egg migration and pathogenesis [35], AChE-KI (or unmutated) eggs were injected into the subserosal layer of the ileum of mice, as previously described [36]. The ileum draining mesenteric lymph node (MLN) cells were isolated from mice on day 5 post injection and divided into two groups to undertake flow cytometric analysis and to quantify secreted cytokines (IFNγ, IL-4 and IL-13) in culture supernatants following 72 hours culture Flow cytometric analysis did not reveal significant difference in the number of CD4+ T cells isolated from mice injected with AChE-KI eggs compared with those injected with eggs electroporated with CON (**Fig 3C**). However, the number of activated CD44hi CD4+ T cells, GATA3+ and IL-4+ T cells were significantly increased in mice injected with eggs treated with X5-KI (**Fig 3**); increased cell numbers of CD4+GATA3 and CD4+IL-4 cells were observed in mice injected with X7-KI eggs compared with mice injected with CON eggs. **S8 Fig** presents the gating strategy for flow cytometric quantification of IFNγ, GATA3, IL-4 and IL-13 levels produced by MLN cells.

The levels of secreted cytokines (IL-4, IL-13 and IFN-γ) were also determined in supernatants of MLN cells after *in vitro* culture for 72 hours. The levels of IL-4 and IL-13 secreted by MLN cells isolated from mice injected with eggs treated with X5-KI were significantly enhanced compared with those from control mice injected with *S. mansoni* CON eggs (**Fig 3C**). In contrast, no change was observed in the level of IFNγ generated by MLN cells isolated from mice injected with treated or untreated eggs (**Fig 3C**).

## Discussion

We demonstrated CRISPR/Cas9-mediated site-specific AChE modification in *S. mansoni* eggs. Given gRNAs with different target sites might guide different endonuclease activities [37, 38], we first pre-screened the activity of two gRNAs (X5 and X7) combined with/without appropriate ssODNs within the AChE loci. Importantly, we found the major modification induced by the CRISPR/Cas9–mediated editing in eggs was HDR. The effect of this modification was further reflected by the *in vivo* immune responses induced by AChE-KI eggs in different tissues of mice, thereby uniquely identifying the phenotypic difference through targeting differing gRNAs.

In our NGS analysis of amplicons from AChE-KI eggs we found, using CRISPResso2 software and a 1 bp window size, that the majority (>95%) of modified reads with NHEJ were due to substitutions, and not insertions or deletions. Furthermore, we determined that the frequency of NHEJ substitutions increased equally across all analysed samples, including control groups, when the window size was changed from 1 bp to the entire amplicon length. This suggests the NHEJ substitutions detected by CRISPResso2 are likely false positives introduced through PCR or sequencing errors although further study is required to confirm these observations. Deletions and insertions were rare in AChE-edited eggs, similar to the report by Sankaranarayanan et al. [16], where CRISPR/Cas9 was used to edit the *SULT-OR* in *S. mansoni* eggs. A possible explanation for these results might be low expression levels of some key NHEJ repair enzymes required for CRISPR/Cas9 editing in schistosome eggs [16], including gene Smp_211060 identified in *S. mansoni* as a homolog of the *KU70/KU80* genes which are essential for the NHEJ pathway [39-41]. The limited NHEJ indels detected in AChE-edited eggs suggested the possibility that most DSBs in AChE loci can be repaired by re-ligating directly without insertion or deletions. However, when ssODN was provided, the DSBs generated by CRISPR/Cas9 editing were repairable by HDR, which is a different outcome to that observed with vertebrate cells where the HDR of the DSBs is extremely low compared to NHEJ, at least where both pathways are equally available [42]. In some single celled parasites including *Plasmodium* [43-46], *Trichomonas vaginalis* [47] and *Cryptosporidium parvum* [48], HDR is the only mechanism of DSB repair due to the lack of the NHEJ pathway; this leads to more accurate repair with less unintended or off-target effects. However, in *Toxoplasma gondii*, frequent NHEJ of DSBs occurs, resulting in random modifications [49, 50]. Precise custom modifications via HDR in *T. gondii* are generated by disrupting the NHEJ pathway through deleting *KU80*, the key gene for NHEJ [39-41], leading to more specific gene editing. Our study shows that the CRISPR/Cas9 mediated editing system in schistosome eggs (delivered by electroporation) was able to induce HDR more prominently; this may provide a mechanism to generate loss-of-function insertions into site-specific nucleases in the schistosome genome and afford a new approach for studying gene editing in blood flukes.

HDR efficiency can be utilized to increase gene ablation [51] reflected by a marked reduction in protein activity. The reduced AChE activity detected in AChE-KI eggs of *S. mansoni*, together with the modified immune response in mice induced by AChE-KI eggs, suggest that the amount of modification (∼0.12 by HDR mediated repair) observed in the genomic NGS analysis, albeit small, nevertheless led to a pronounced phenotypic change at the protein level in this multicellular parasite. The other recent CRISPR/Cas9 studies on parasitic flatworms [52], including targeting *ω1* in *S. mansoni* [15] and *Ov-grn-1* in *O. viverrini* [17], reported similar findings. Schistosomes are complex, multi-cellular, multi-organ invertebrates that have evolved a range of adaptations to survive in diverse environments, and these features present considerable challenges for successful genomic editing. Incomplete modification of the AChE gene may stimulate *S. mansoni* to alter cholinergic signalling transduction and further regulate downstream signalling pathways (such as the PI3K and ERK pathways), thereby amplifying the effect of the knockdown. In addition, as reported in CRISPR/Cas9 editing in *S. mansoni* [16], *Strongyloides* [53] and *C. elegans* [54], large deletions might be missed by amplicon sequencing. Undetected modification(s) during AChE editing may have led to the pronounced phenotypic changes observed at the protein level in these helminths but this needs to be further explored.

The low efficacy of CRISPR/Cas9 mediated gene editing of AChE achieved in *S. mansoni* eggs may be due to particular characteristics of this multicellular parasite now considered.

1. The schistosome egg has a protective eggshell with a hardened structure comprising three layers [55] that may make it difficult to penetrate [56]. Nevertheless, targeting the egg may provide optimal access to schistosome germ line cells in order to introduce insertional mutagenesis in chromosomes of the blood fluke [57]; this is because the egg has a relatively simple structure and a high stem-like cell content [58] compared with other life cycle stages,
2. Using a pool of several thousands of liver eggs for CRISPR/Cas9-mediated editing may result in low knockdown efficacy. Liver eggs represent a mixture of eggs of variable age and different stages of development and in *S. mansoni* higher transgene efficiency at the genomic level occurs in immature compared with mature eggs [57]. The use of freshly laid eggs by adult schistosomes cultured *in vitro* may improve on the knockdown effect but the logistics involved in collecting sufficient numbers of these immature eggs for downstream sequencing analysis and phenotype studies present a major challenge.
3. Variation in the pattern of CRISPR/Cas9 induced modifications may depend on the function or distribution of the targeted gene. The AChE protein is located throughout different components of schistosome eggs [26] and is highly expressed in mature eggs (**S9 Fig**) in which the miracidia have developed within the eggshell into multi-cellular, mobile, ciliated larvae composed of organs, tissues, muscles and nerves. CRISPR/Cas9-mediated AChE editing in the nuclei of cells within the thin cellular epithelia of individual mature eggs may have been more frequent than in cells deeper within the egg including the extra-embryonic inner envelope. When AChE-KI eggs were injected into mice, the edited AChE distributed in the egg epithelia would be readily released from the eggs into the surrounding tissues thereby modulating host immune responses.

In this study, our NGS analysis focused on AChE-edited eggs. However, as reported by Sankaranarayanan *et al* [16], adult *S. mansoni* might be a better stage to achieve higher efficiency of CRISPR/Cas9 mediated modifications compared with eggs (and sporocysts). Accordingly, an improved programmed genome editing may occur in AChE-edited adults, which we now plan to investigate.

Schistosome eggs play critical roles in host pathogenesis, immune modulation, and the transmission of schistosomiasis. Eggs trapped in the intestinal wall or liver can induce a strong Th2 immune response which is associated with the chronic pathology characteristic of schistosomiasis [59]. A novel mouse infection model, involving intravenous injection of eggs via the tail vein, revealed enhanced Th2 responses in lung cells and splenocytes isolated from mice injected with X5-KI eggs. The role of *S. mansoni* X5-KI eggs in suppressing the Th2 immune response was further confirmed when intestinal injection of these mutated eggs was performed, a superior method for assessing the generation of specific anti-egg immune responses. MLNs isolated from the intestine, where eggs are locally injected, provide a more sensitive readout of generated immune responses and the approach more closely resembles the natural site of egg transition and granuloma formation [35]. The MLN analysis indicated that knockdown of AChE targeting gRNA X5 in eggs could trigger a superior host immune response than that targeting gRNA X7, although NGS analysis of X5-KI and X7-KI eggs showed similar levels of HDR modifications (∼ 0.12%). The upstream location of gRNA X5 in the genome compared with X7 allowed insertion of stop codons delivered by X5ssODN into the DSB of X5 generated by CRISPR/Cas9 editing, thereby affecting the expression of downstream DNA sequences of AChE. This further suggests that motifs or domains of AChE that are involved in regulating the Th2 immune response might be located between the X5 and X7 sites in the AChE gene.

The important role of cholinergic signalling in a helminth parasite influencing the host immune system was recently reported in a study where *Trypanosoma musculi* was engineered to heterogeneously express a secreted AChE from *Nippostrongylus brasiliensis;* the secreted protein was shown to alter the cytokine environment by enhancing production of IFNγ and TNFα, with a concomitant reduction in IL-4, IL-5 and IL-13 levels [60]. We also previously showed a predominantly Th1-type protective immune response characterized by increased production of IFNγ in mice vaccinated with recombinant AChE [26] following schistosome challenge, indicating egg-secreted AChE might be critical in inhibiting the host IL-4 response. These prior studies support the findings reported here where X5-KI eggs elicited an increased Th2 response characterized by elevated levels of IL-4, IL-5, IL-10 and IL-13 secreted by splenocytes. These cytokines are predominantly produced by Th2 cells in various models of type 2 immunity [61, 62] and they inhibit Th1 responses [63]. This may account for the reason why we did not observe any changes in the levels of IL-2 and IFNγ which are characteristically produced by Th1 cells [64]. To confirm *in vivo* alveolar immune responses induced by the AChE-KI eggs in mice, lung homogenates were used in qPCR assays to determine the mRNA expression of IL-4, INFγ, Ctla4 and RetnlA. The level of IL-4 was increased only in mice injected with X5-KI eggs complementing the significantly increased level of IL-4 observed in the lung cells and splenocytes. The expression of Ctla4, which inhibits the Th2 response [65], was increased in mice injected with AChE-X7 KI eggs, and this was reflected by the increased level of TNF (typifying a Th1 response) observed in splenocytes isolated from mice injected with AChE-X7 KI eggs. RetnlA is known to be involved in the down modulation of Th2 responses [66] and is expressed by alternatively activated macrophages [66]. However, there was no significant difference observed in the RetnlA expression in all groups, indicating that AChE-KI in eggs may be associated with Ctla4-mediated Th2 suppression, or that the effects were too localised around the granulomas to allow their detection.

Using the murine i.v. injection model we showed a strong IFNγ response was evident at 2-weeks post injection in all mice across the different groups analysed. At this same time point, a significant decrease in lung granuloma size was observed in all the mice injected with KI eggs. However, an enhanced Th2 response was evident only in mice injected with X5-KI eggs indicating the size of granulomas may keep increasing, driven by the climbing Th2 response around the X5-KI eggs, but this needs further investigation.

The circumoval granulomas represent an accumulation of host immune cells including macrophages, eosinophils and neutrophils, around schistosome eggs [67]. The host immune cells act to protect the surrounding host tissue from toxins produced by the egg, resulting in a physical barrier and the sequestering of these secreted egg products [68]; yet these cells are ineffective at clearing infection [69]. To date, the underlying mechanisms of granuloma formation remain poorly understood, and our current knowledge is based on *in vitro* studies using cells that have not been obtained from granulomas, thereby hindering an improved understanding of the dual functions of immune killing and healing of granulomas generated by schistosome eggs [69]. Granuloma formation is driven by chemokines released by T lymphocytes and resident liver cells, including HSC, which attract migrating immune cells to the site of egg deposition [70]. Inhibition of HSC activation can lead to decreased collagen production and fibrosis formation [30, 71], features also evident in this current study. We found the growth of HSCs (LX-2 cells) was significantly inhibited when the cells were incubated with SEA from AChE-KI eggs compared with SEA extracted from unmutated eggs. Given activated HSCs are recruited to developing granulomas [30], a possible explanation is that the inactivation or inhibition of HSCs by the SEA of AChE-KI eggs may have resulted in the migration of fewer immune cells (macrophages, eosinophils and neutrophils) and a decline in the lungs in the expression of collagen type 1-α1 (COL1α1), a marker of collagen deposition [72], around the granuloma. The combination of these features *in vivo* might directly lead to the formation of small granulomas. Our results are supported by previous studies showing paeoniflorin (a monoterpene glycoside), which was shown to down-regulate the activity of AChE in rats [73], reduced fibrosis in mice infected with schistosomes as well as inhibiting HSC proliferation and collagen synthesis [74]. However, whether there are key differences between the way lung and hepatic granulomas develop remains unclear. Further investigation is required to improve our understanding of the modulation of granuloma formation induced by schistosome eggs entrapped in lungs including the dynamic changes in the host response correlating with secreted cytokines and development and subsequent resolution of the granuloma.

### Concluding comments

Establishment of the CRISPR/Cas9-mediated KI system in *S. mansoni* makes it feasible to generate specific loss-of-function modifications in this parasitic flatworm. Notwithstanding the progress presented here, and in earlier reports on CRISPR/Cas9 activity in *S. mansoni* [15, 16], future improvement in the efficiency of gene editing can be anticipated from additional modifications and refinements of the methodology. This genome editing approach can be readily extended to the study of other schistosome genes and to those of other trematode species thereby paving the way for high-throughput functional analysis of flatworm genes generally.

## Methods

### Maintenance of *S. mansoni*

All experiments were approved by the Animal Ethics Committee of the QIMR Berghofer Medical Research Institute and performed in quarantine-approved facilities. The study was conducted according to the guidelines of the National Health and Medical Research Council of Australia, as published in the Australian Code of Practice for the Care and Use of Animals for Scientific Purposes, 7th edition, 2004 (www.nhmrc.gov.au). All work related to live *S. mansoni* life cycle stages was performed in quarantine-accredited premises.

Swiss mice (females, 6 weeks old) were infected with 100 *S. mansoni* cercariae. Seven weeks post-infection mice were euthanised and adult worms were obtained by portal perfusion with 37°C DMEM medium (Invitrogen, Carlsbad, USA). Adult worms were incubated in complete schistosome medium (CSM) containing DMEM medium, supplemented with 10% (v/v) heat-inactivated fetal calf serum, 100 IU/ml penicillin and 100 μg/ml streptomycin, at 37°C in an atmosphere of 5% CO2 in air overnight. Mouse livers were removed at necropsy and eggs were isolated and purified as described [75]. The eggs were cultured in the CSM at 37°C under 5% CO2 in air. Soluble worm antigen preparation (SWAP) and soluble egg antigen (SEA) were prepared from adult worms and eggs as described [76, 77].

### Guide RNA target selection and reconstruction of GeneArt CRISPR Nuclease Vector

Single guide RNA (gRNA) target sequences (<15 bp from the desired edit) were selected using the web-based tools available for CRISPR design prediction programs: (1) http://bioinfogp.cnb.csic.es/tools/breakingcas/ [78], and (2) Benchling Referral Program (https://benchling.com) to predict cleavage sites for the *Streptococcus pyogenes* Cas9 nuclease within the genome of *S. mansoni*. The gRNA targeted exon 5 and 7 of the AChE (Gene ID Smp_154600, www.genedb.org), residues 722-741 (named X5), 1738-1757 (named X7), respectively, adjacent to the protospacer adjacent motif, AGG (Fig 1A). The Smp_154600 includes 9 exons interspersed with 8 introns (57.78kb) (Fig 1A). A double stranded DNA sequence complementary to the gRNA was inserted into GeneArt CRISPR Nuclease Vector (Life technologies, Carlsbad, USA) according to the manufacturer’s instructions, which encodes Cas9 from *Streptococcus pyogenes* driven by the human cytomegalovirus (CMV) immediate early promoter and sgRNA driven by the human U6 promoter. The sequences of the two gRNA of Smp_154600 are shown: X5-gRNA, CACCAGGTAATATGGGTCTC (Fig 1B); X7-gRNA, TGGGCTAACTTTGCACGCAC (Fig 1C) Control vector was reconstructed using negative non-function gRNA (GACCAGGATGGGCACCACCC). Those gRNAs were synthesized by Integrated DNA Technologies (Singapore).

Two single-stranded oligodeoxynucleotides (ssODN) targeting AChEX5 (X5ssODN), AChEX7 (X7ssODN) were synthesized by Integrated DNA Technologies (Singapore). The sequences of the three ssODNs are shown: X5ssODN, with homology arms of 50 nt each in length at the 3’ (position 738-787 nt) and 5’ (688-737 nt) flanks and a small transgene (5’-<u>TAA</u>G<u>TGA</u>C<u>TAG</u>G<u>TAA</u>C<u>TGA</u>G<u>TAG</u>C-3’, encoding stop codons (six) in all reading frames) (Fig 1B); X7ssODN, with homology arms of 50 nt each in length at the 3’ (position 1754-1803nt) and 5’(1704-1753nt) flanks and a small transgene (5’-<u>TAA</u>G<u>TGA</u>C<u>TAG</u>G<u>TAA</u>C<u>TGA</u>G<u>TAG</u>C-3’, encoding stop codons (six) in all reading frames) (Fig 1C);

### Transfection of adult *S. mansoni* and eggs with a CRISPR Vector with/without ssODN

Pools of 10,000 of eggs or 10 pairs of adult *S. mansoni* were subjected to transfection by electroporation in 100 µl Opti-MEM containing: 1) Medium only (Medium); 2) X5-gRNA and X5ssODN (3 µg for each) (X5-KI); 3) X5-gRNA (3 µg) (X5); 4) X7-gRNA and X7ssODN (3 µg for each) (X7-KI); 5) X7-gRNA (3 µg) (X7); 6) X5-gRNA, X5ssODN and X7-gRNA, X7ssODN (3 µg for each) (X5+X7-KI); 7) X5-gRNA and X7-gRNA (3 µg for each) (X5+X7); 8) GeneArt® CRISPR Nuclease Vector reconstructed with negative gRNA (3 µg) (CON); 9) X5ssODN (3 µg); 10) X7ssODN (3 µg). The mixture was pipetted into a 4 mm pre-chilled electroporation cuvette and subjected to a square wave with a single 20 ms impulse at 125 v [79] (Gene Pulser Xcell Electroporator, Bio-Rad, USA), and subsequently maintained at 37°C, 5% CO2 in air for 2 days. The eggs and adults were collected on day 2 post electroporation and genomic DNA was extracted from eggs and adults using E.Z.N.A. SQ Tissue DNA Kit (OMEGA bio-tek, Norcross, USA).

The five controls included parasites subjected to electroporation in the presence of medium only (medium), GeneArt® CRISPR Vector reconstructed with negative gRNA (CON), ssODN only (X5ssODN and X7ssODN). Wide type parasites were cultured without electroporation (WT).

### PCR amplification to detect knock out/in into exon 5 and exon 7 of AChE

Targeting gRNA-AChEX5, PCR assays were performed on each genomic DNA samples extracted from eggs and adults using distinct primer pairs (S1 Table). The primer pairs CF+R4 and F2+R4, which amplify locations 22,607-23,411 nt and 22,982-23,411 nt of Smp_154600, respectively, served as a positive control for the presence of genomic DNA with the Smp<u>_</u>154600 of AChE. The other primer pairs CF+R and F2+R shared one reverse primer (R) complementary to the knock-in 124 nt transgene with two forward primers, CF, F2 at positions 22,607-22,626 and 22,982-23,006 nt, respectively, bind complementary to two sites of Smp<u>_</u>154600 targeting site gRNA-AChEX5 DSB (Fig 1B).

Targeting gRNA-AChEX7, PCR assays were carried out on genomic DNA samples extracted from eggs and adults using distinct primer pairs (S1 Table). The primer pairs F1+R1 and F1+R6, to amplify locations 37,932-38,518 nt and 37,932-38,403 nt of Smp_154600, respectively, were used as a positive control for the presence of genomic DNA with the Smp<u>_</u>154600. The other primer pairs F+R1 and F+R6 shared one forward primer (F) complementary to the knock-in 124 nt transgene with two reverse primers, R1, R6 at positions 38,499-38,518 and 38,384-38,403 nt, respectively (Fig 2A).

The PCR mix included 10 µl Green GoTaq DNA polymerase mix (Promega, Madison, USA) with 200 nM of each primer and 10 ng genomic DNA. Thermal cycling conditions involved denaturation at 95°C, 3 min followed by 30 cycles of 94°C, 30 sec, 60°C, 30 sec and 72°C, 30 sec and a final extension at 72°C for 5 min. Amplicons were visualized following agarose gel electrophoresis (1.2% agarose/TAE), and those of the expected sizes were extracted and purified from gels for sequencing to confirm the presence and knock-out/in of the transgene. Sanger sequencing analysis of the KI amplicons was used to confirm the insertion of the transgene into the AChE locus at X5 and X7 at the predicted cleavage sites.

### Illumina sequencing

Pooled egg genomic DNA samples from 11 experimental groups of AChEX5 or X7 with/without ssODN were used as the template to amplify the on-target DNA fragment using MiSeq primers (including paired primers F2+R, F2+R4, F+R6, F1+R6) (Fig 1D) with NEBNext Ultra II Q5 Master Mix (New England Biolabs, Ipswich, USA). PCR reactions were performed with 60 ng DNA samples from different experimental groups in 50 µl reaction mix using the PCR program 98°C for 30sec of denaturation followed by 35 cycles of 98°C for 10 sec, 65°C for 30 sec, 72°C for 30 sec and final extension at 72°C for 2 min. The amplicons of expected sizes were purified by using QIAquick Gel Extraction kit (Qiagen, Hilden, Germany). Amplicons generated from 2-4 different PCR reactions from each group were pooled, and 100ng of amplicons from each sample were used to construct the uniquely indexed paired-end read libraries (BGI, Hong Kong). These libraries were pooled and the pooled library was quantified using a bioanalyzer (Agilent 2100, Agilent Technologies, Santa Clara, USA) combined with StepOnePlus Real-Time PCR System (Applied Biosystems™, Carlsbad, USA) to measure the adapters before sequencing. The qualified libraries were sequenced on MiSeq using paired-end 300bp reads (Illumina, San Diego, USA). In total, 22 multiplexed samples (S2 Table) were run on MiSeq.

Amplicon NGS libraries are available at the European Nucleotide Archive under the study accession number EGAS00001004455.

### CRISPResso2 analysis

To detect the amount of CRISPR/Cas9-mediated editing in *S. mansoni* eggs, raw fastq files from individual samples were run through the command line version of CRISPResso2 (v2.0.30) [28]. The following parameters were used for CRISPResso2 analysis: “fastq_r1”, “fastq_r2”, “amplicon_seq”, “guide_seq”, “expected_hdr_amplicon_seq”, “trimmomatic_options_string” to provide Illumina adapters, “max_paired_end_reads_overlap” of 200, “exclude_bp_from_left” equal to the length of the respective forward primer + 1, and “exclude_bp_from_right” equal to the length of the respective reverse primer + 1. The remaining parameters were kept at default values. Notably, the “window_around_sgrna” parameter was kept at 1 to ensure PCR and/or sequencing artefacts were not counted as false positive NHEJ insertions, deletions, or substitutions. Separately, we tested increasing values of the “window_around_sgrna” parameter (20, 100, and 0, where 0 disables the argument to consider the entire length of the amplicon) to determine its effect on the occurrence of NHEJ and HDR events.

To confirm potential HDR reads reported by CRISPResso2, we parsed the “Alleles_frequency_table.txt” output file using a custom python script. First, to ensure each read matched the respective forward and reverse primers, the 5’ end of the read (i.e. start of the read to the length of the forward primer) was compared to the sequence to the forward primer, and the 3’ end of the read (i.e. end of the read to the length of the reverse primer) was compared to the reverse complement of the reverse primer, and allowed for 1 bp mismatch using the ‘levenshtein’ function within the ‘distance’ python package. These filtered reads were then used to determine the editing frequencies of NHEJ and HDR. The HDR reads reported by CRISPResso2 were then confirmed using fasta36 (v3.8; https://github.com/wrpearson/fasta36), where an HDR read was considered ‘confirmed’ if it had > 90% identity to the expected knock-in sequence.

### AChE activity assays

Cultured adult worms and eggs were collected on day 2 after electroporation for protein extraction of soluble worm antigen preparation (SWAP) [76] and soluble egg antigen (SEA) [32] as described. The AChE activity in these SWAP (7.5 μg/mL) and SEA (0.45 μg/mL) preparations were measured using the Amplex Red Acetylcholine/Acetylcholinesterase Assay Kit (Invitrogen) according to the manufacturer’s instructions.

### *In vitro* scratch wound assays

*In vitro* scratch wound assays were carried out to determine the effect of SEA extracted from control and CRISPR/Cas9 mutated eggs [80, 81]. HSC cells (LX-2) were cultured in completed medium containing high glucose EMDM (Invitrogen), 2% fetal bovine serum (FBS), 1% glutamax and 1% Pen/strep. Cells were grown to create a confluent monolayer in 96-well plates.

Then the monolayers were scraped in a straight line to create a “scratch” using the Wound Maker-IncuCyte ZOOM-Image Lock Plate system (Essen Bioscience, Michigan, USA) and washed once with the culture medium. Cells were continued to culture in the fresh completed medium containing 30 µg/ ml SEA. Residual endotoxin was assessed by using an Endotoxin Standards kit (Lonza, Anaheim, USA) to ensure there was no LPS contamination in the SEA extracted from control and AChE-KI eggs). The growth of cells was monitored by IncuCyte Zoom for 3 days. Images were acquired for each well at 3 hours interval by an in-built phase contrast microscope. The area of the scratch was measured using ImageJ (image processing program, Java). The total area (μm^2^) was obtained at every time point until would closure, and triplicate measures were compared to calculate an average closure rate per group (area of original wound/time to closure in hours). All data were then statistically analysed using GraphPad Prism Software V7.

### Pathological changes and immune responses induced by intravenous injection of AChE-KI eggs in mice

Mutated and unmutated egg batches were intravenously (i.v.) injected into mice under sterile conditions as described [82]. AChE-KI eggs were cultured for 2 days following electroporation with mixtures of X5-KI; X7-KI; X5+X7-KI, respectively. Briefly, 1,000 eggs in 100 µl of sterile PBS were injected into the lateral tail vein of female Swiss mice (7–8 weeks of age). Mice in control groups were injected with PBS (naive), untreated eggs (WT) or with eggs electroporated with negative control GeneArt vector (Con). All mice were euthanised 2 weeks after injection. This experiment was repeated twice with each group comprising 5 mice.

### Measurement of lung granuloma size in mice

For each mouse, the left lung was fixed in 4% (v/v) formalin and paraffin-embedded sections of these samples were prepared and stained with Haematoxylin and Eosin (H&E). Slides were digitized using an Aperio Slide Scanner (Aperio Technologies, Vista, USA). The degree of lung pathology was quantified by measurement of the area density of granulomatous lesions using Aperio Image Scope v11.1.2.760 software (Leica Biosystems Imaging, Buffalo Grove, USA), and was estimated from the area of the granulomas divided by the total area of the liver tissue in the image.

### Immune responses induced by intravenous injection of AChE-KI eggs

1. IgE response determination. Bloods were collected from each mouse at 2 weeks after injection, and sera were prepared and stored at −80°C. The total IgE level of individual sera was measured using an IgE mouse ELISA kit (Thermo Fisher Scientific, Waltham, USA), according to the manufacturer’s instructions.
2. Cytokine analysis. The spleen and lung of each mouse (*n* = 5 per group) were collected and splenocytes [83] and lung cells [84] were isolated as described. Briefly, the individual lung was finely chopped and digested in 1mL of lung digestion buffer, containing 4mg/mL collagenase D (Sigma-Aldrich, St. Louis, USA), 10% FCS (Fetal Calf Serum, Invitrogen) 0.5mg/mL DNase (Promega) in IMDM (Iscoves Modified Dulbecco’s Medium, Invitrogen), for 1hr at 37°C and 150 RPM in a shaking incubator. The digested lung was then mechanically disrupted through a syringe followed by filtering through a 70µm cell strainer. The individual spleen was pressed through a 70µm cell strainer. All the spleen and lung samples were then lysed using RBC lysis buffer (Sigma-Aldrich, Missouri, USA) to remove red blood cells. Following repeated washing and centrifugation steps, the cells were resuspended in IMDM containing 10% (v/v) fetal bovine serum, 100 μg/ml Streptomycin, 100 Units/ml Penicillin and 0.05 mM 2-Mercaptoethanol. Cell counts were determined by trypan blue exclusion, and then cells were seeded into 96-well plates (5 × 10^5^ cells/well). The cells were incubated in the presence of SEA (2 μg/well, extracted from *S. mansoni* liver eggs) for 72 hours at 37°C. Stimulation with ConA (0.1 μg/well) was performed as a positive control and non-stimulated splenocytes or lung cells were used as a negative control. Each cell culture supernatant (50µl/well) was harvested and secreted cytokines (IFNγ, TNFα, IL-2, 4, 5, 6, 10, 13) were assayed using LEGEND plex Mouse Th1/Th2 panel kits (BioLegend, San Diego, USA) according to the manufacturer’s instructions.
3. Real time PCR. A small loop of lung was cut individually from each mouse injected i.v. with AChE-KI eggs. Total RNA was extracted from the lung tissue using Trizol (Invitrogen) and an RNeasy Mini Kit (Qiagen) [85] followed by DNase (Promega) digestion; cDNA was synthesised as described [86]. Real-time PCR was performed and analysed using the ABI ViiA™ 7 real time PCR system (Thermo Fisher Scientific). TaqMan Gene Expression Assays (ThermoFisher) for IL-4 (Mm00445259_m1); INFγ (Mm01168134_m1), Ctla4 (Mm00486849) and Retnla (resistin-like molecules, Mm00445109) were employed to detect potential disturbances in Th2 polarization. HPRT (hypoxanthine guanine phosphoribosyl transferase) was used as a reference gene to normalise all data generated [87].

The expression level of COL1α1 in lung tissue of mice injected with mutated eggs was determined by qPCR quantification using SYBR Green master mix (Applied Biosystems, Foster City, USA) on a Corbett Rotor Gene 6000 (Corbett Life Sciences, Uithoorn, Netherlands). cDNA obtained from lung tissue of mice in treated and untreated groups was used as template and primer sequences for COL1α1 were as previously reported [88, 89]. Data were analysed by importing the standard curve to each run using Rotor-Gene 6000 Series software (version 1.7).

### Immune responses induced by intestinal injection of AChE-KI eggs into mouse lymph nodes

Egg intestinal injection surgery. Laparotomy surgery was performed as described [36]. In brief, mice were anaesthetised using Isoflurane (Abbot Animal Health), a small incision into the skin and the muscle layer of the animal’s midline was made using a scalpel and extended using scissors. The intestine was carefully displayed onto a surgical cloth using cotton buds and moistened. Mutated or unmutated *S. mansoni* eggs were resuspended at 4,000 eggs/50µl PBS and 4,000 eggs were injected into the subserosal layer of the ileum at two locations using Micro-Fine Plus Hypodermic Syringes (29G x 12.7mm; BD Bioscience). After successful egg injection, the intestines were replaced into the body cavity and the muscle and skin incisions closed using 6.0 Vicryl absorbable sutures (Johnson and Johnson). The analgesics Buprenorphine (0.1 mg/kg; Vetergesic, Reckitt Benckiser Healthcare) and Carprofen (5 mg/kg; Rimadyl, Pfizer) were administered subcutaneously into the flank and mice were closely monitored post-surgery to ensure full recovery from the anaesthesia and monitored on a daily basis thereafter. This experiment was repeated twice, with each group comprising 4-6 mice.

Cell isolation and *in vitro* stimulation. Mice were scarified on day 5 post egg injection and the ileum draining mesenteric lymph nodes were isolated as described [36]. Single cell suspensions were obtained by disrupting each lymph node with a 70µm cell strainer (BD Bioscience). In each experiment, cells were split to perform flow cytometric analysis and to set up re-stimulation cultures from the same sample. 2 x 10^6^ cells were incubated in RPMI 1640 (Life Technologies), supplemented with 2.5 ng/ml PMA (Sigma-Aldrich), 1 mg/ml ionomycin (Invitrogen), 0.5% GolgiStop (BD Bioscience, Franklin, USA) and 10% FCS for four hours at 37 °C, after which live/dead staining was performed and cell surface markers were stained. Cells were fixed and permeabilized using the eBioscience Foxp3/Transcription Factor Staining Buffer Set (eBioscience, San Diego, USA) and intracellular staining was performed following the manufacturer’s instructions. For re-stimulation cultures, 1 x 10^6^ MLN cells were cultured in X-vivo 15 medium (Lonza) supplemented with 1% L-glutamine (Invitrogen), 0.1% 2-mercaptoethanol (Sigma-Aldrich) and 7.5 µg/ml SEA in round bottom 96-well plates (Corning) at 37°C and 5% CO2. Supernatants were collected after three days and cytokine levels were determined using IL-4, IL-13 and IFN-γ “ready-set-go” ELISA kits or paired antibodies (eBioscience).

Antibodies for flow cytometric analysis. The following combination of fluorescently labelled primary antibodies against cell surface markers, intracellular cytokines and transcription factors were used: anti-CD3 (17A2), anti-CD4 (clones GK1.5), anti-CD8a (53–6.7), anti-CD44 (IM7), anti-CD45R/B220 (RA3-6B2) and anti-IL4 (11B11) from Biolegend, and anti-IFN-γ (XMG1.2) and anti-GATA3 (TWAJ) from eBioscience. Cells were analysed using a BD LSRFortessa flow cytometer running FACSDiva Software (BD Bioscience) and analysed using FlowJo Software (Tree Star).

### Statistical analysis

All data are presented as the mean ± SE. Differences between groups were assessed for statistical significance using One-way ANOVA; Tukey post hoc test or Kruskal–Wallis were used for comparisons involving > 2 groups compared with the control. GraphPad Prism software (Version 7, GraphPad Software, La Jolla, CA, USA) was used for all statistical analyses. *P* values ≤ 0.05 were considered to be statistically significant. \**p*<0.05, \*\**p*<0.01, \*\*\**p*<0.001; not significant (ns).

## Supplementary information

**S1 Fig**. Discrete exon/intron structure of the loci encoding genes Smp_136690 and Smp_125350 (two paralogues of Smp_154600, acetylcholinesterase) in *S. mansoni*.

**S2 Fig**. Amino acid alignment of proteins Smp_136690, Smp_125350 and Smp_154600. Red boxes indicate the two conserved subdomains including the carboxylesterase type-B signature 2 (E163-P173 in Smp_154600) and carboxylesterase type-B serine active site (F264-G279 in Smp_154600). Motifs include N-myristoylation sites boxed in purple (G77-L82, G101-Q106, G304-N309, G401-E406, G539-Y544 in Smp_154600). The conserved active catalytic triad site (shown as a red star) is located at E406 in Smp_154600, while the 4 residues (W154, W192, Y208, Y544 in Smp_154600, in red triangles) in the rings of 14 aromatic amino acid residues of *S. mansoni* AChE, which contribute to the 3D-structure of AChE, are conserved in the appropriate locations in the two molecules.

**S3 Fig. (A**) PCR products visualized in ethidium bromide-stained agarose gels demonstrating Cas9-catalyzed target site-specific insertional mutagenesis in exon 5 and 7 of the AChE gene. Genomic DNA extracted from adult *S. mansoni* treated with CON, X5-KI, X5, X7-KI, X7 and X5+X7-KI, respectively, was used as template for PCR. Evidence for transgene knocked-in into the programmed target site revealed by amplicons of the expected sizes in lanes CFR and FR1 of 682 and 392 bp, respectively; these span the mutated site in the genomic DNAs pooled from adult *S. mansoni*, including positive controls flanking the insert site of gRNA-X5 (CFR4, 805 bp) and the insert site of gRNA-X7 (F1R1, 587 bp). The control DNA was isolated from adult worms electroporated with medium only. (B) AChE activity of soluble worm antigen preparation (SWAP) extracted from adult *S. mansoni* that had been treated with medium, CON, X5-KI, X5, X7-KI, X7 and X5+X7-KI, respectively. SWAP extracted wild type (WT) adult worms were used as positive control.

**S4 Fig**. Electroporation of eggs does not induce a significant difference in egg viability. Eggs isolated from the liver of an *S. mansoni*-infected mouse were hatched. (A) The hatching rates were calculated before and after electroporation. Data are presented as the mean ± SE; t-test analysis showed *p*=0.16. (B) Hatched miracidia visualised under the microscope.(a) mature egg in the process of hatching; (b) miracidium escaping from an egg; (c) empty eggshell after hatching; (d) free-swimming miracidium in medium following hatching; (e) immature egg. Scale bars, 100µm.

**S5 Fig. Effect of increasing the CRISPResso2 window parameter on the quantification of modified reads**. Stacked bar plots of the number of modified reads expressed as a percentage of aligned reads per experimental sample. The CRISPResso2 “-w” window parameter was changed to a (A) value of 1, (B) value of 20, (C) value of 100, and (D) value of 0 (the entire amplicon length) while keeping all other CRISPResso2 parameters constant. The results of each of the four analyses have been separated into two groups: experimental samples amplified using primers targeting AChE Exon 5 (upper panels) and experimental samples amplified using primers targeting AChE Exon 7 (lower panels).

**S6 Fig** Images of the growth of LX-2 cells on day 0, 1 and day 2 after incubation with soluble egg antigens (SEA) extracted from: (A) wild type (WT) eggs; (B) eggs electroporated with medium; (C) CON eggs; (D) X5-KI eggs; (E) X7-KI eggs and (F) X5+X7-KI eggs. Scale bars, 300μm.

**S7 Fig. Cytokine mRNA expression observed in the lungs of mice i**.**v. injected with CRISPR/Cas9 mutated/unmutated eggs**. A portion of mouse lung homogenate was used to determine, by qPCR, the expression levels of: (A) IL-4, (B) INFγ, (C) RetnlA (resistin-like molecules) and (D) Ctla4. One-way ANOVA analysis was used to establish statistical significance compared with the CON:* *p*<0.05, \*\**p*<0.01, \*\*\**p*<0.001.

**S8 Fig. Gating strategy for the assessment of Th1 and Th2 cells in the MLN after subserosal injection with *S. mansoni* eggs**. MLNs were harvested 5 days after subserosal egg injection, digested and stimulated with PMA/ionomycin for 4 hours. CD4 T cells were identified by flow cytometry analysis by gating on live single cells that were B220-CD3+ and CD4+. IFN-γ+ CD4+ Th1 cells and GATA3+ and IL-4+ Th2 cells were quantified after the injection of WT *S. mansoni* eggs or eggs treated with negative control vector (CON), X5-KI, X7-KI or X5-KI+X7-KI.

**S9 Fig. Immunolocalisation showing that the expression of AChE by *Schistosoma* eggs in liver tissue is distributed throughout the eggs [26] and is highly expressed in mature eggs**. Adapted from our previous work [26], we showed that AChE is located in the thin cellular epithelium, the extra-embryonic inner envelope of eggs, and in cells of the inner-most cellular layers of advanced *S. japonicum* egg granulomas [26]. AChE is highly expressed in the mature egg compared with immature egg, in the former of which the miracidium has developed within the eggshell into a multi-cellular, mobile, ciliated larva comprising organs, tissues, muscles and nerves [26]. Given the high level of conservation (88% amino acid identity) in protein sequences for AChE in *S. japonicum* and *S. mansoni* [25], we assume the distribution of AChE in the eggs of these two species will be similar. IM, immature egg; M, mature egg. Scale bars, 100µm

**S1 Table. Sequences of guide RNA and primers used for PCR amplification**

**S2 Table. Quantification of the modifications in NGS sequence reads using CRISPResso2 software**.

## Acknowledgements

*B. glabrata* snails provided by the NIAID Schistosomiasis Resource Center of the Biomedical Research Institute (Rockville, MD) through NIH-NIAID Contract HHSN272201700014I for distribution through BEI Resources. This work received support from an Australian Infectious Disease Research Centre Seed Grant and a Program Grant from the National Health and Medical Research Council of Australia (APP 1037304) and from a Strategic Award from the Wellcome Trust, award no. 107475/Z/15/Z, entitled the Flatworm Functional Genomics Initiative (KF Hoffmann, PI, PJ Brindley, co-PI).

## Author Contributions

### Conceptualization

Hong You, Haran Sivakumaran, Patrick Driguez, Juliet D. French, Wannaporn Ittiprasert, Paul J. Brindley, Donald P. McManus

### Data curation

Hong You, Johannes U. Mayer, Rebecca L. Johnston, Lambros T. Koufariotis, Nicola Waddell

### Formal analysis

Hong You, Johannes U. Mayer, Rebecca L. Johnston, Olga Kondrashova, Lambros T. Koufariotis, Nicola Waddell

### Funding acquisition

Hong You, Malcolm K. Jones, Donald P. McManus

### Investigation

Hong You, Juliet D. French, Nicola Waddell, Paul J. Brindley, Malcolm K. Jones, Donald P. McManus

### Methodology

Hong You, Johannes U. Mayer, Rebecca L. Johnston, Haran Sivakumaran, Shiwanthi Ranasinghe, Vanessa Rivera, Xiaofeng Du, Mary G. Duke, Wannaporn Ittiprasert

### Project administration

Hong You, Malcolm K. Jones, Donald P. McManus

### Resources

Hong You, Mary G. Duke, Donald P. McManus

### Supervision

Hong You, Paul J. Brindley, Malcolm K. Jones, Donald P. McManus

### Validation

Hong You, Johannes U. Mayer, Haran Sivakumaran, Juliet D. French, Nicola Waddell, Paul J. Brindley, Malcolm K. Jones, Donald P. McManus

### Writing – original draft

Hong You, Johannes U. Mayer, Rebecca L. Johnston, Olga Kondrashova, Haran Sivakumaran

### Writing – review & editing

Hong You, Juliet D. French, Nicola Waddell, Paul J. Brindley, Malcolm K. Jones, Donald P. McManus

